# Toll-9 interacts with Toll-1 to mediate a feedback loop during apoptosis-induced proliferation in *Drosophila*

**DOI:** 10.1101/2020.10.19.346254

**Authors:** Alicia Shields, Alla Amcheslavsky, Elizabeth Brown, Yingchao Nie, Takahiro Tanji, Y. Tony Ip, Andreas Bergmann

## Abstract

*Drosophila* Toll-1 and all mammalian Toll-like receptors regulate innate immunity. However, the functions of the remaining eight Toll-related proteins in *Drosophila* are not fully understood. Here, we show that *Drosophila* Toll-9 is necessary and sufficient for a special form of compensatory proliferation after apoptotic cell loss (undead apoptosis-induced proliferation (AiP)). Mechanistically, for AiP, Toll-9 interacts with Toll-1 to activate the intracellular Toll-1 pathway for nuclear translocation of the NF-κB-like transcription factor Dorsal which induces expression of the pro-apoptotic genes *reaper* and *hid*. This activity contributes to the feedback amplification loop that operates in undead cells. Given that Toll-9 also functions in loser cells during cell competition, we define a general role of Toll-9 in cellular stress situations leading to the expression of pro-apoptotic genes which trigger apoptosis and apoptosis-induced processes such as AiP. This work identifies conceptual similarities between cell competition and AiP.

## Introduction

Since the discovery of the original *Toll* gene in *Drosophila* (*Toll-1* hereafter), a large number of Toll-related genes have been identified in both insects and mammals (reviewed in (Anthoney et al., 2018; Bilak et al., 2003)). The *Drosophila* genome encodes a total of 9 Toll-related proteins including Toll-1 (Ooi et al., 2002; Tauszig et al., 2000), while mammalian genomes encode between 10 to 13 Toll-like receptors (TLR). TLRs are single-pass transmembrane proteins which upon ligand-stimulation usually trigger a conserved intracellular signaling pathway, culminating in the activation of NF-κB transcription factors (Anthoney et al., 2018).

Initially identified as an essential gene for dorsoventral patterning in the early *Drosophila* embryo (Anderson et al., 1985a; Anderson et al., 1985b), *Toll-1* was later also found to be an essential component for innate immunity (Lemaitre, 2004; Lemaitre et al., 1996). In this function, Toll-1 signaling via the NF-κB transcription factors Dorsal and Dorsal-related immunity factor (Dif) induces the expression of anti-microbial peptides (AMPs) which mediate innate immunity (Meng et al., 1999; Rutschmann et al., 2000; Tauszig et al., 2000). A role in innate immunity has also been demonstrated for all mammalian TLRs (Lemaitre, 2004). Likewise, *Drosophila* Toll-8 (aka Tollo) regulates immunity in the trachea (Akhouayri et al., 2011). In addition, *Toll-7* may regulate anti-viral responses, albeit through an NF-κB-independent mechanism (Lamiable et al., 2016; Nakamoto et al., 2012). However, for the remaining Toll-related proteins in *Drosophila*, a function in innate immunity has not been clearly demonstrated (Anthoney et al., 2018; Tauszig et al., 2000).

Of particular interest is Toll-9 in *Drosophila* because it behaves genetically most similar to *Toll-1* (Bettencourt et al., 2004; Ooi et al., 2002) and its intracellular TIR domain is most closely related to those of the mammalian TLRs (Khush and Lemaitre, 2000; Ooi et al., 2002; Wasserman, 2000). Over-expression of *Toll-9* results in the production of AMPs which led to the proposal that Toll-9 might be involved in innate immunity (Ooi et al., 2002). However, a loss-of-function analysis with a defined *Toll-9* null allele did not confirm this prediction (Narbonne-Reveau et al., 2011). Nevertheless, although *Drosophila* Toll-9 does not appear to be directly involved in regulating expression of AMPs, it has been implicated together with a few other Toll-related proteins in cell competition, an organismal surveillance program which monitors cellular fitness and eliminates cells of reduced fitness (losers) (Alpar et al., 2018; Meyer et al., 2014). Depending on the type of cell competition, Toll-9 participates in the expression of the pro-apoptotic genes *reaper* and *hid* in loser cells triggering their elimination (Meyer et al., 2014). Therefore, it has been proposed that the function of Toll signaling for elimination of bacterial pathogens by AMPs and for elimination of unfit cells by pro-apoptotic genes bears a conceptual resemblance between innate immunity and cell competition (Alpar et al., 2018; Meyer et al., 2014).

Apoptosis is an evolutionarily well conserved process of cellular suicide mediated by a highly specialized class of Cys-proteases, termed caspases (reviewed by (Fuchs and Steller, 2011; Salvesen et al., 2016; Shalini et al., 2015)). In *Drosophila*, the pro-apoptotic genes *reaper* and *hid* promote the activation of the initiator caspase Dronc (Caspase-9 homolog in *Drosophila)*. Dronc activates the effector caspases DrICE and Dcp-1 (Caspase-3 and −7 homologs) which trigger the death of the cell (reviewed in (Salvesen et al., 2016; Shalini et al., 2015)). However, caspases not only induce apoptosis, but they can also have non-apoptotic functions (reviewed by (Aram et al., 2017; Baena-Lopez, 2018)) such as apoptosis-induced proliferation (AiP) during which caspases in apoptotic cells promote the proliferation of surviving cells independently of their role in apoptosis (Diwanji and Bergmann, 2019; Fogarty and Bergmann, 2017).

Early work has shown that AiP requires the initiator caspase Dronc (Fan et al., 2014; Huh et al., 2004; Kondo et al., 2006; Ryoo et al., 2004; Wells et al., 2006). To reveal the mechanism of AiP, we are expressing the pro-apoptotic gene *hid* and the apoptosis inhibitor *p35* simultaneously using the *ey-Gal4* driver (referred to as *ey>hid,p35*) which drives expression in the larval eye disc anterior to the morphogenetic furrow (MF) (Fan et al., 2014). *p35* encodes a specific inhibitor of the effector caspases DrICE and Dcp-1, but does not block the activity of Dronc (Meier et al., 2000; Yoo et al., 2002). In *ey>hid,p35-expressing* discs, the apoptotic pathway is activated by *hid* expression, but blocked downstream due to DrICE and Dcp-1 inhibition by P35, rendering cells in an “undead” condition. While apoptosis is inhibited in undead tissue, AiP still occurs due to non-apoptotic signaling by the initiator caspase Dronc, triggering hyperplasia of the anterior portion of the larval eye imaginal discs at the expense of the posterior eye field which is reduced in size (Fan et al., 2014). Combined, these effects result in overgrowth of the adult head capsule (Figure 1B), but a reduction or even absence of the eyes in the adult animal (Fan et al., 2014).

**Figure 1.**
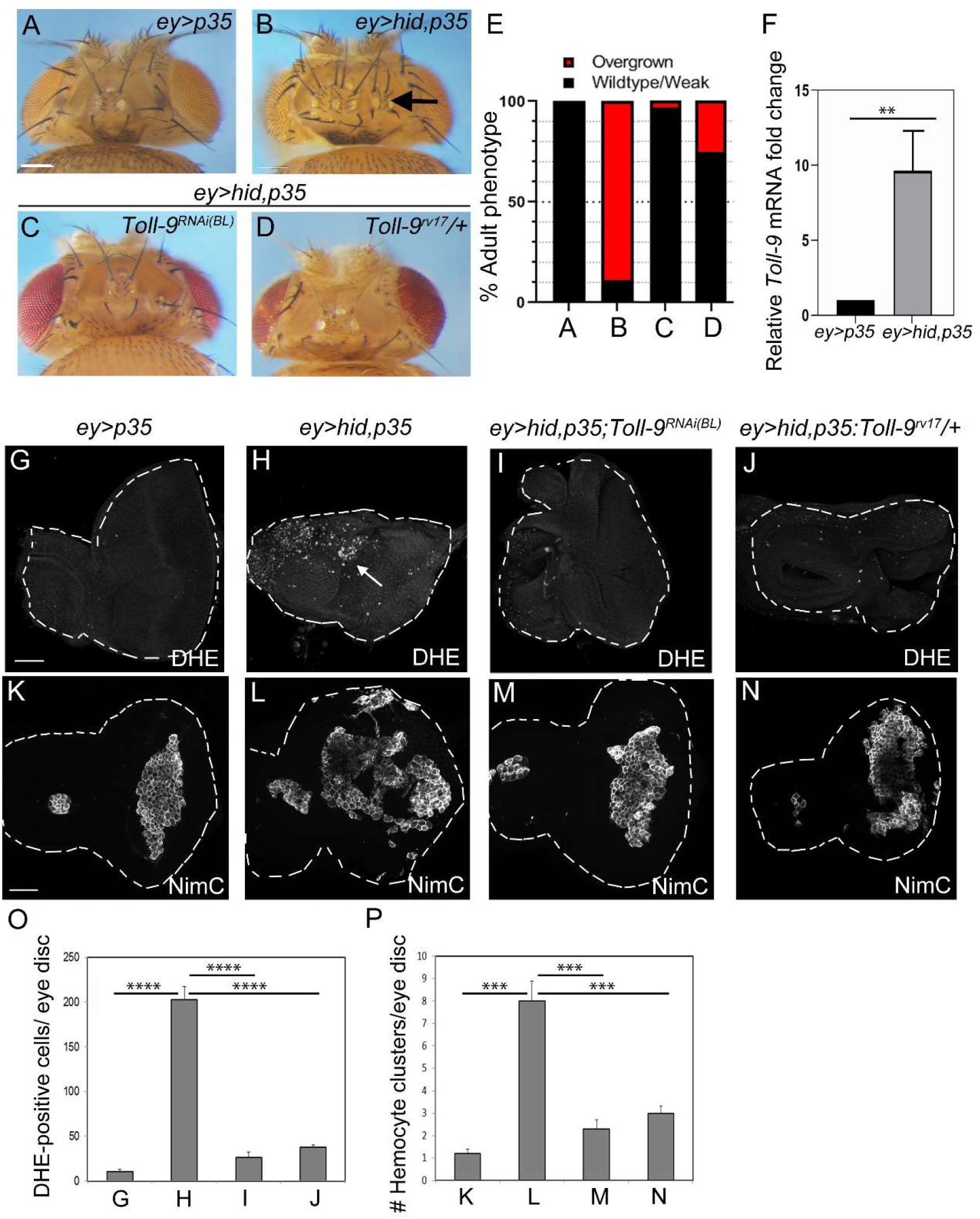
*Toll-9* is required for *ey>hid,p35-induced* overgrowth, ROS generation and hemocyte activation. The *Toll-9^RNAi(BL)^* line in (B, H, L) is Bloomington stock #BL35035.(**A-D**) Reduction of *Toll-9* activity by RNAi (C) or the heterozygous *Toll-9^rv17^* mutant (D) suppresses *ey>hid,p35-induced* overgrowth (B). Arrow in (B) points to additional bristles and ocelli. Expression of *p35* (*ey>p35*) (A) does not trigger overgrowth. Scale bar: 200μm. (**E**) Quantification of the data in (A-D). Based on qualitative screening criteria, overgrowth of progeny is scored by the presence of additional bristles and ocelli, expansion of the head capsule and amorphic overgrowth. In severe cases, eye tissue is lost. At least 100 flies were scored for each genotype. (**F**) *ey>hid,p35* induces *Toll-9* expression. Relative mRNA levels of *Toll-9* in *ey-Gal4* and *ey>hid,p35* genetic backgrounds. The average of three independent experiments is plotted using Student’s t-test, one-tailored distribution, two sample unequal variance. **P<0.01. Plotted is the mean intensity ± SEM. (**G-N**) ROS generation (G-J) and hemocyte number (K-N) in *ey>hid,p35* eye imaginal discs are strongly reduced by *Toll-9* RNAi and the heterozygous *Toll-9^rv17^* mutant. Dihydroethidium (DHE) was used as ROS indicator. Arrow in (H) highlights DHE labeling. Anti-NimC antibody was used as hemocyte marker. Scale bars: 50μm. (**O,P**) Quantification of the data in (G-J) and (K-N), respectively. DHE counts and hemocyte clusters were counted across entire discs. Data were analyzed by one-way ANOVA with Holm-Sidak test for multiple comparisons. Plotted is the mean ± SEM. ****p <0.0001; ***P<0.001. The following numbers of discs were analyzed: 4 (G), 8 (H), 4 (I), 7 (J), 5 (K), 6 (L), 6 (M), 5 (N).

Using the undead model, we showed that AiP is mediated by extracellular reactive oxygen species (ROS) generated by the NADPH oxidase Duox (Fogarty et al., 2016). ROS trigger the recruitment of hemocytes, *Drosophila* immune cells most similar to mammalian macrophages, to undead imaginal discs (Diwanji and Bergmann, 2020). Hemocytes in turn release signals which stimulate JNK activity in undead cells which then promotes AiP (Fogarty et al., 2016). In addition, JNK also induces the expression of *hid* (Moreno et al., 2002), thus setting up an amplification loop in undead cells which continuously signals for AiP (Fogarty et al., 2016).

Given that undead cells are abnormal cells (Martin et al., 2009) with potentially altered cellular fitness and that signaling by Toll-related proteins surveilles cellular fitness (Alpar et al., 2018; Meyer et al., 2014), we examined if signaling by Toll-9 has an important function for undead AiP. Here, we show that *Toll-9* is strongly upregulated in undead cells and is necessary for the overgrowth of undead tissue. Over-expression of *Toll-9* induces all markers of undead AiP signaling including Duox-dependent ROS generation, hemocyte recruitment and JNK signaling. Mechanistically, we provide genetic evidence for a heterologous interaction between Toll-9 and Toll-1 which engages the canonical intracellular Toll-1 signaling pathway to promote nuclear translocation of the NF-κB-like transcription factor Dorsal which induces the expression of *reaper* and *hid*. This activity contributes to the establishment of a feedback amplification loop that signals continuously for AiP. In conclusion, while Toll-9 does not appear to have a function in innate immunity, it appears to be involved in stress situations such as cell competition and AiP which result in the expression of pro-apoptotic genes such as *reaper* and *hid*.

## Results

### Toll-9 is required for overgrowth of undead tissue

We examined if *Drosophila Toll-9* is involved in undead AiP. Indeed, depletion of *Toll-9* by RNAi suppressed *ey>hid,p35-induced* overgrowth (Figure 1A-C; quantified in Figure 1E). We also generated a *Toll-9* deletion mutant by imprecise excision of an existing P-element insertion, referred to as *Toll-9^rv17^* (Supplemental Figure S1A). Heterozygosity of *Toll-9^rv17^* suppressed *ey>hid,p35-induced* overgrowth of undead tissue (Figure 1D,E). These results suggest that *Toll-9* is required for *ey>hid,p35-induced* AiP. Furthermore, by qRT-PCR, we found that *Toll-9* mRNA levels increased 8-10 fold in *ey>hid,p35* undead tissue compared to control tissue (Figure 1F).

### Toll-9 is required for ROS generation and hemocyte recruitment

To place *Toll-9* into the AiP network, we performed a number of genetic tests in the *ey>hid,p35* genetic background. *Toll-9* RNAi suppressed the ectopic JNK and Wg activity in *ey>hid,p35* background suggesting that it acts upstream of these genes (Supplemental Figure S2). Furthermore, *Toll-9* RNAi or *Toll-9^rv17^* heterozygosity strongly suppressed the generation of ROS (Figure 1G-J; quantified in 1O). The same set of experiments with a different ROS indicator, H_2_DCF-DA, gave similar results (Supplemental Figure S3). In addition, hemocytes that require ROS to spread out across most of the *ey>hid,p35* imaginal disc (Diwanji and Bergmann, 2020; Fogarty et al., 2016) displayed the naïve morphology seen at control discs in response to *Toll-9* RNAi or *Toll-9^rv17^* heterozygosity (Figure 1K-N; quantified in 1P). We conclude that *Toll-9* is required for ROS generation and the subsequent recruitment of hemocytes in *ey>hid,p35*-expressing tissue.

### *Toll-9* overexpression in *ey>p35* background engages a similar AiP pathway as *ey>hid,p35*

While *ey>Gal4*-induced expression of *Toll-9* does not induce any visible phenotype (Supplemental Figure S1B,C), co-expression of *Toll-9* with *p35* (*ey>p35,Toll-9*) is sufficient to cause overgrowth of the resulting head cuticle (Figure 2A,B; quantified in 2M). It is unknown why overexpression of *Toll-9* can induce overgrowth only in the presence of *p35* (see Discussion), but we have made a similar observation when overexpressing another factor involved in AiP, called Myo1D (Amcheslavsky et al., 2018). Nevertheless, the *ey>p35,Toll-9*-induced overgrowth phenotype gives us an assay to examine the mechanism by which Toll-9 works in the AiP pathway.

**Figure 2.**
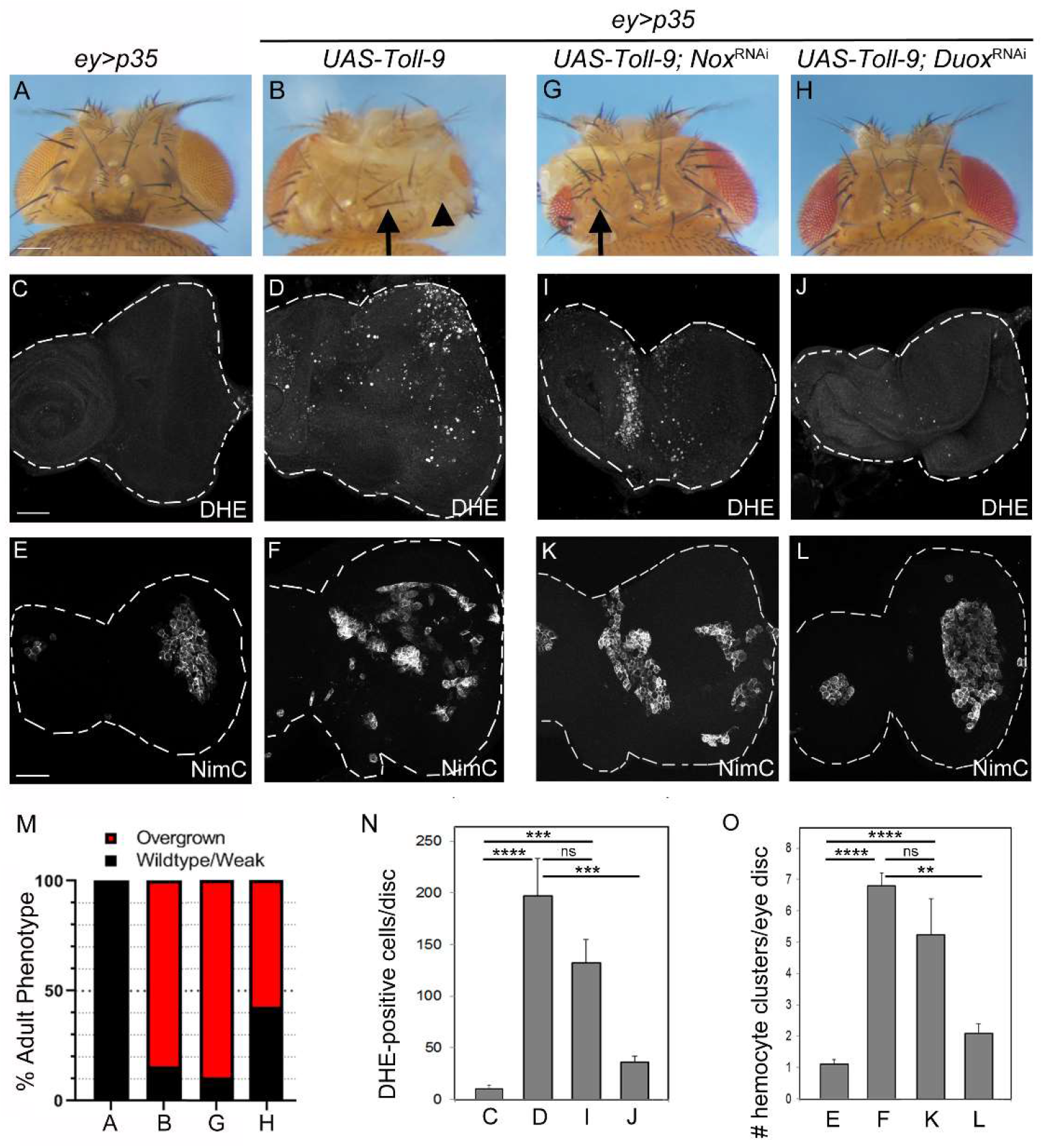
Toll-9 promotes overgrowth of *ey>p35* head capsules and hemocyte activation via Duox-generated ROS. The genotypes of heads and eye discs are indicated above the panels. (**A,B**) Transgenic expression of *Toll-9* causes overgrowth of the head capsule of *ey>p35-* animals. Quantified in (M). The arrow in (B) highlights additional bristles, the arrowhead overgrown cuticle at the expense of eye tissue. Scale bar: 200μm. (**C-F**) Transgenic expression of *Toll-9* in *ey>p35* eye imaginal discs triggers ROS generation (D) and hemocyte recruitment (F). DHE and NimC were used as ROS indicator and hemocyte marker, respectively. Quantified in (N,O). Scale bars: 100μm. (**G,H**) *Duox* RNAi (H), but not *Nox* RNAi (G), can suppress *Toll-9-induced* overgrowth of *ey>p35* head capsules. Quantified in (M). The arrow in (G) highlights overgrown tissue with additional bristles. (**I-L**) ROS generation and hemocyte clustering of *ey>p35,Toll-9* eye discs are suppressed by *Duox* RNAi (J,L), but not by *Nox* RNAi (I,K). DHE and NimC were used as ROS indicator and hemocyte marker, respectively. Quantified in (N,O). (**M**) Quantification of the data in (A,B,G,H). Scoring criteria as in Figure 1E. At least 100 flies were scored for each genotype. (**N,O**) Quantification of the DHE counts in (C,D,I,H) and hemocyte clustering in (E,F,K.L). DHE counts and hemocyte clusters were counted across entire discs. Data were analyzed by one-way ANOVA with Holm-Sidak test for multiple comparisons. Plotted is the mean ± SEM. ****p <0.0001; ***P<0.001; **P<0.01. ns - not significant. n=: 4 (C), 4 (D), 7 (E), 12(F), 6 (G), 6 (H), 6 (I), 6 (J), 7 (K), 6 (L).

The expression of *Toll-9* in *ey>p35* discs is associated with the generation of ROS as well as recruitment of hemocytes (Figure 2C-F; quantified in Figure 2N,O), consistent with the loss of ROS and hemocyte recruitment by *Toll-9* depletion in *ey>hid,p35* background (Figure 1I,J,M,N). We showed previously that the membrane-bound NADPH oxidase Duox is required for ROS generation in *ey>hid,p35* imaginal discs, while the second NADPH oxidase, Nox, is not (Fogarty et al., 2016). Consistently, *Duox* RNAi, but not *Nox* RNAi, suppressed the head overgrowth, the generation of ROS, and the recruitment of hemocytes in eye imaginal discs of *ey>p35, Toll-9* animals (Figure 2G,H,I,J,K,L; quantified in Figure 2M-O) suggesting that Toll-9’s overgrowth-promoting effect through ROS generation by Duox, but not Nox, is required for the recruitment of hemocytes at *ey>p35,Toll-9* imaginal discs.

JNK signaling is known to mediate signaling and overgrowth in the undead *ey>hid,p35* AiP pathway (Fan et al., 2014; Ryoo et al., 2004). Consistently, expression of a RNAi transgene against JNK (*bsk^RNAi^*) suppressed the head overgrowth of *ey>p35, Toll-9* adult animals (Supplemental Figure S4A-C). We also observed an increase of JNK activity in *ey>p35, Toll-9* imaginal discs based on the JNK reporter *TRE-RFP* (Supplemental Figure S4D-G) suggesting that JNK is induced and required for Toll-9-induced overgrowth in the *ey>35* background.

These similarities between *ey>hid,p35* and *ey>p35, Toll-9* imply that overexpression of *Toll-9* in *ey>35* background engages a similar hyperplasia pathway compared to *ey>hid,p35* animals.

However, in *ey>hid,p35* discs, *hid* induces caspase, i.e. Dronc activity, which is required for AiP (Fan et al., 2014; Huh et al., 2004; Kondo et al., 2006; Wells et al., 2006). Therefore, we examined if *ey>p35, Toll-9* induces caspase activity. For that purpose, we used the cleaved Dcp-1 (cDcp-1) antibody which is commonly used as a marker of caspase activity in *Drosophila* tissue (Li et al., 2019). Indeed, while there is no significant increase of caspase activity in *ey>Toll-9* discs, there is increased cDcp-1 labeling evident in *ey>p35,Toll-9* imaginal eye discs (Figure 3C–E; quantified in 3F). Furthermore, depletion of *dronc* by RNAi resulted in strong suppression of *ey>p35, Toll-9* induced overgrowth (Figure 3A,B) suggesting that *Toll-9* overexpression can induce Dronc activity in the presence of *p35*. These data raise the question of how Toll-9 can achieve this activity.

**Figure 3.**
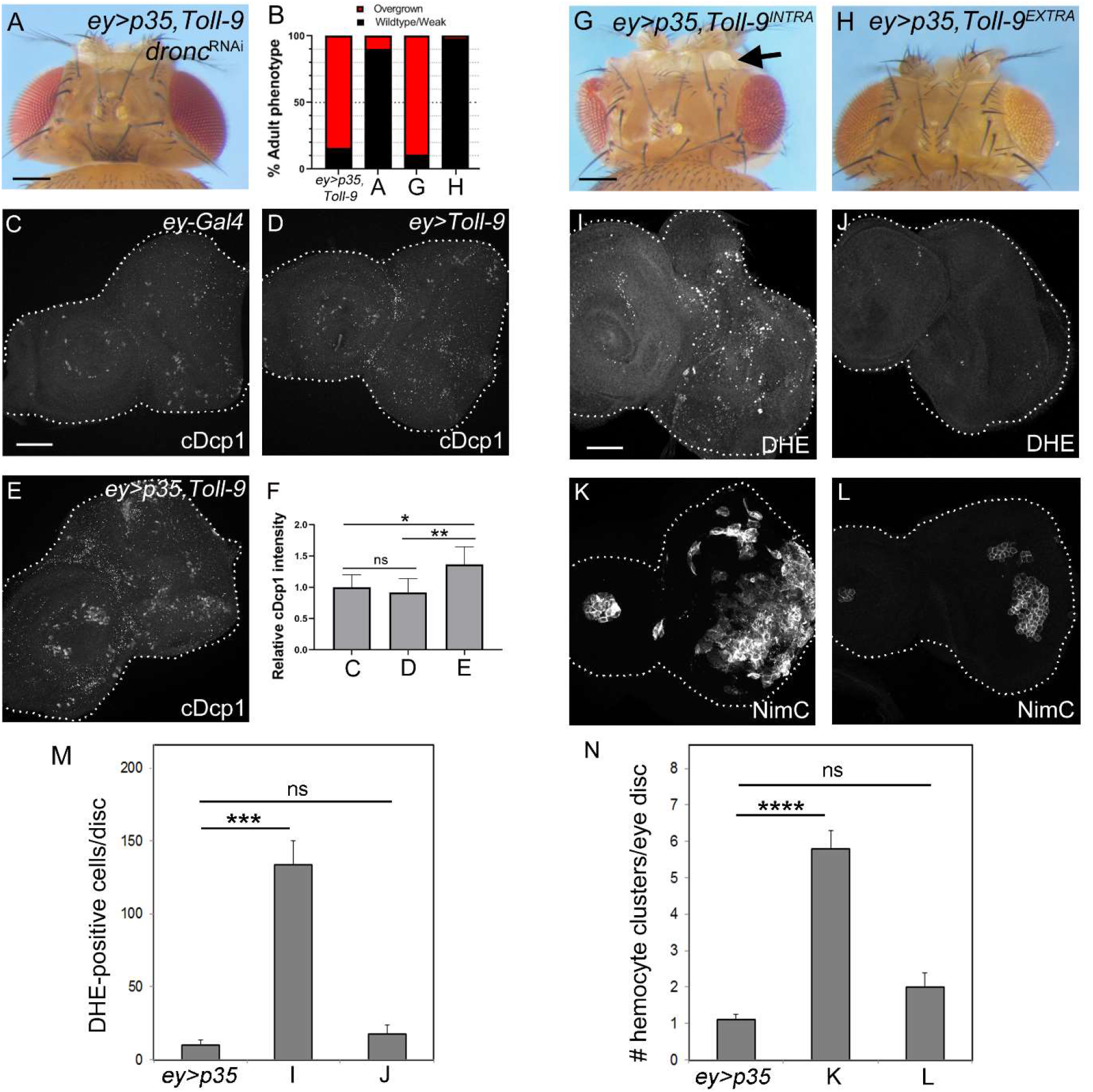
The intracellular domain of Toll-9 is sufficient for overgrowth of *ey>p35-* expressing head capsules, ROS generation and hemocyte recruitment. (**A**) RNAi-knockdown of *dronc* suppressed *ey>p35, Toll-9-induced* overgrowth. Quantified in (B). Scale bar: 200μm. (**B**) Quantification of the data in (A,G,H). Scoring criteria as in Figure 1E. At least 100 flies were scored for each genotype. (**C-E**) cDcp1 labeling of imaginal eye discs of *ey-Gal4* (C) *ey>Toll-9* (D) and *ey>p35,Toll-9* (E) third instar larvae. Quantified in (F). Scale bar: 100μm. (**F**) Quantification of the data in (C-E). cDcp1 labeling was measured across entire discs and normalized to disc size. Data were analyzed by one-way ANOVA with Holm-Sidak test for multiple comparisons. Relative signal intensity is plotted ± SEM. **P<0.01; *P<0.05. ns - not significant. n=: 7 (C), 9 (D), 12 (E). (**G,H**) Transgenic expression of the intracellular domain of *Toll-9* (*Toll-9^INTRA^*) (G), but not the extracellular domain (*Toll9^EXTRA^*) (H), causes overgrowth of the head capsule of *ey>p35*-expressing animals. Quantified in (M). The arrow in (G) highlights overgrown amorphic tissue. Scale bar: 200μm. (**I-L**) Transgenic expression of *Toll-9^INTRA^*, but not *Toll-9^EXTRA^*, triggers ROS generation (I,J) and hemocyte clustering (K,L) in *ey>p35*-expressing eye imaginal discs. DHE and NimC were used as ROS indicator and hemocyte marker, respectively. Quantified in (M,N). Scale bar: 100μm. (**M,N**) Quantification of the data in (I,J) and (K,L). DHE counts and hemocyte clusters were obtained across entire discs. Data were analyzed by one-way ANOVA with Holm-Sidak test for multiple comparisons. Plotted is the mean ± SEM. ****p <0.0001; ***P<0.001. ns - not significant. The following numbers of discs were analyzed: 4 (*ey>p35* in M), 6 (I), 7 (J), 5 (*ey>p35* in N), 10 (K), 4 (L).

### Toll-9 requires the intracellular domain to induce overgrowth

To address this question, we determined first whether Toll-9 mediates this activity in the AiP network through an intracellular signaling pathway or through cell-cell adhesion mediated by the extracellular domain (Anthoney et al., 2018; Keith and Gay, 1990; Ward et al., 2015). We separately expressed intra- and extracellular domains of Toll-9 (*Toll-9^INTRA^* and *Toll-9^EXTRA^*, respectively) linked to the signal peptide and transmembrane domain, in *ey*>*p35* background. Expression of *Toll-9^INTRA^* is sufficient to induce overgrowth of *ey>p35* head capsules, accompanied by ROS generation and hemocyte activation (Figure 3G,I,K; quantified in Figure 3B,M,N). In contrast, expression of *Toll-9^EXTRA^* did not trigger these phenotypes (Figure 3H,J,L; quantified in Figure 3B,M,N). These data suggest that the intracellular, but not the extracellular, domain of Toll-9 is sufficient for the generation of ROS, hemocyte activation and overgrowth of *ey*>*p35*-expressing tissue, most likely by activating an intracellular signaling pathway.

To explore the possibility that Toll-9 engages an NF-κB-like transcription factor for this overgrowth, we tested the two NF-κB factors involved in Toll-1 signaling in *Drosophila*, Dorsal and Dif (Ip et al., 1993; Meng et al., 1999; Rutschmann et al., 2000; Steward, 1987). Consistently, RNAi targeting either *dorsal* or *Dif* suppressed the overgrowth of *ey>p35, Toll-9* expressing animals (Figure 4A-C; quantified in Figure 4H). Further support for the notion of an intracellular pathway comes from the observation that knockdown of *Myd88* and a mutant of *tube*, known intracellular components acting upstream of Dorsal and Dif in the Toll-1 pathway (Horng and Medzhitov, 2001; Letsou et al., 1991; Tauszig-Delamasure et al., 2002), also suppressed *ey>p35, Toll-9-induced* overgrowth strongly and moderately, respectively (Figure 4D,E; quantified in Figure 4H). RNAi against another gene involved downstream in Toll-1 signaling,*pelle* (Shelton and Wasserman, 1993; Towb et al., 2001), weakly suppressed *ey>p35,Toll-9* (Figure 4F,H).

**Figure 4.**
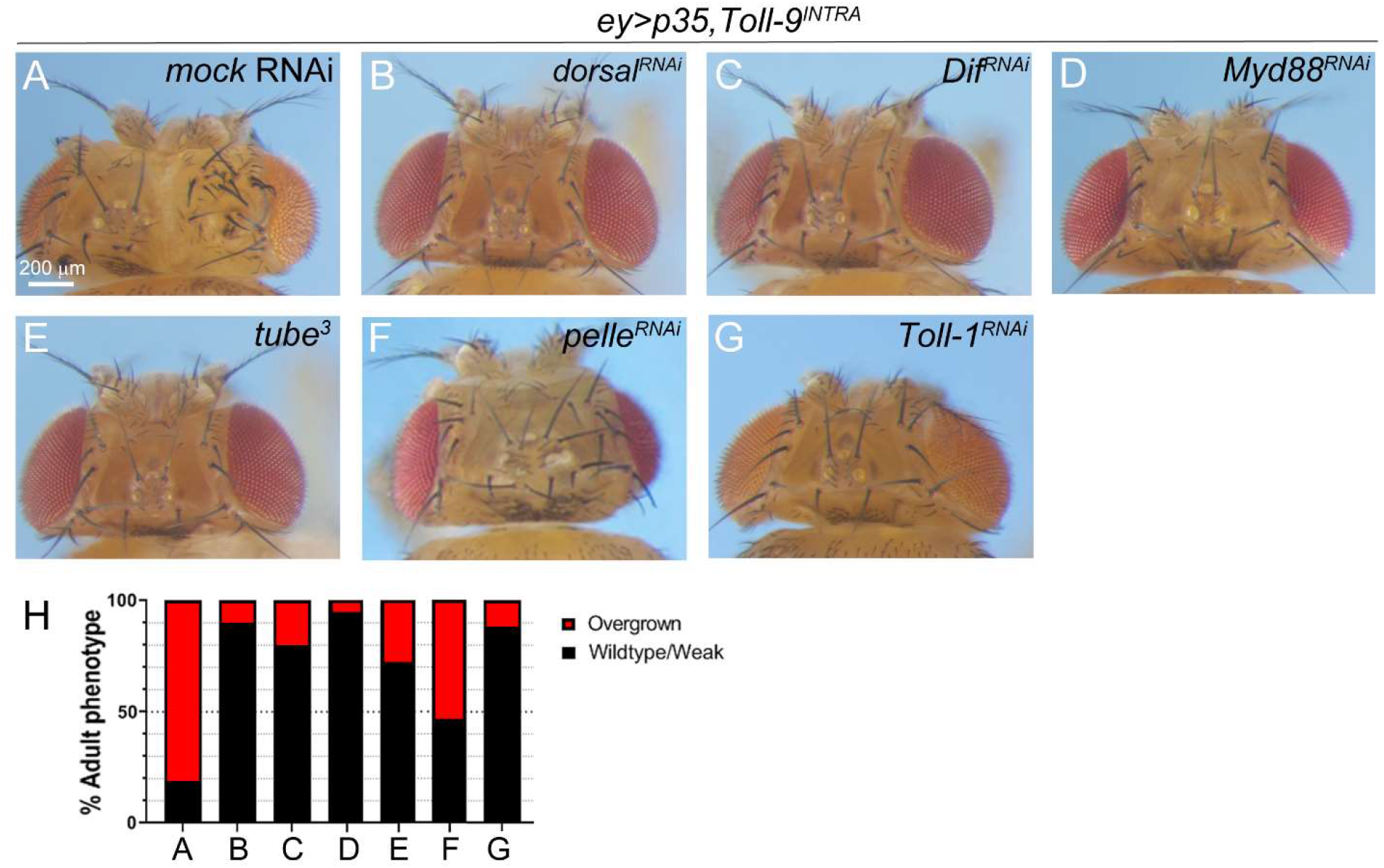
Canonical, intracellular Toll-1 signaling contributes to *ey>p35,Toll-9-induced* overgrowth. (**A-C**) *ey>p35,Toll-9-induced* overgrowth is strongly suppressed by RNAi-mediated knockdown of *dorsal* (B) and *Dif* (C). Scale bar: 200μm. (**D-G**) RNAi-mediated knockdown of *Myd88* (D), *pelle* (F) and *Toll-1* (G) as well as a *tube* mutant (E) suppressed *ey>p35, Toll-9-induced* overgrowth. (**H**) Quantification of the data in panels (A-G). Scoring criteria as in Figure 1E. At least 100 flies were scored for each genotype.

One function of the intracellular Myd88/Tube/Pelle pathway is to mediate the nuclear translocation of the NF-κB transcription factor Dorsal (Roth et al., 1989; Rushlow et al., 1989; Steward, 1989). Therefore, we examined if misexpression of Toll-9 can trigger nuclear localization of Dorsal in wing imaginal discs using *ptc-Gal4* as a driver. We chose *ptc-Gal4* in these experiments, because the striped expression of *Toll-9* in the *ptc* domain allows side-by-side comparisons between *ptc*-expressing and *ptc*-non-expressing areas. Consistent with the expectation, there is strong nuclear accumulation of Dorsal in the *ptc* domain upon *Toll-9* misexpression compared to control (Figure 5A,B; see also orthogonal sections on top of the panels). Interestingly, while misexpression of *Toll-9* alone (*ptc>Toll-9*) is sufficient to trigger nuclear translocation of Dorsal independently of *p35* (Figure 5B^2^), in the presence of *p35* (*ptc>p35,Toll-9*), the expression domain of *ptc* is strongly enlarged (compare Figure 5C^4^ with 3B^4^) further supporting the notion that expression of *Toll-9* induces hyperplasia only in the presence of *p35* (see Discussion). To determine if the intracellular Toll pathway is required for the nuclear translocation of Dorsal, we examined if knockdown of *Myd88* which is the strongest suppressor of *ey>p35,Toll-9-induced* overgrowth (Figure 4H), can block the nuclear localization of Dorsal. Indeed, depletion of *Myd88* by RNAi resulted in strong loss of nuclear localization of Dorsal (Figure 5D). Combined, these data suggest that Toll-9 overexpression engages the intracellular Toll-1 pathway involving Myd88, Tube and Pelle for nuclear localization of Dorsal.

**Figure 5.**
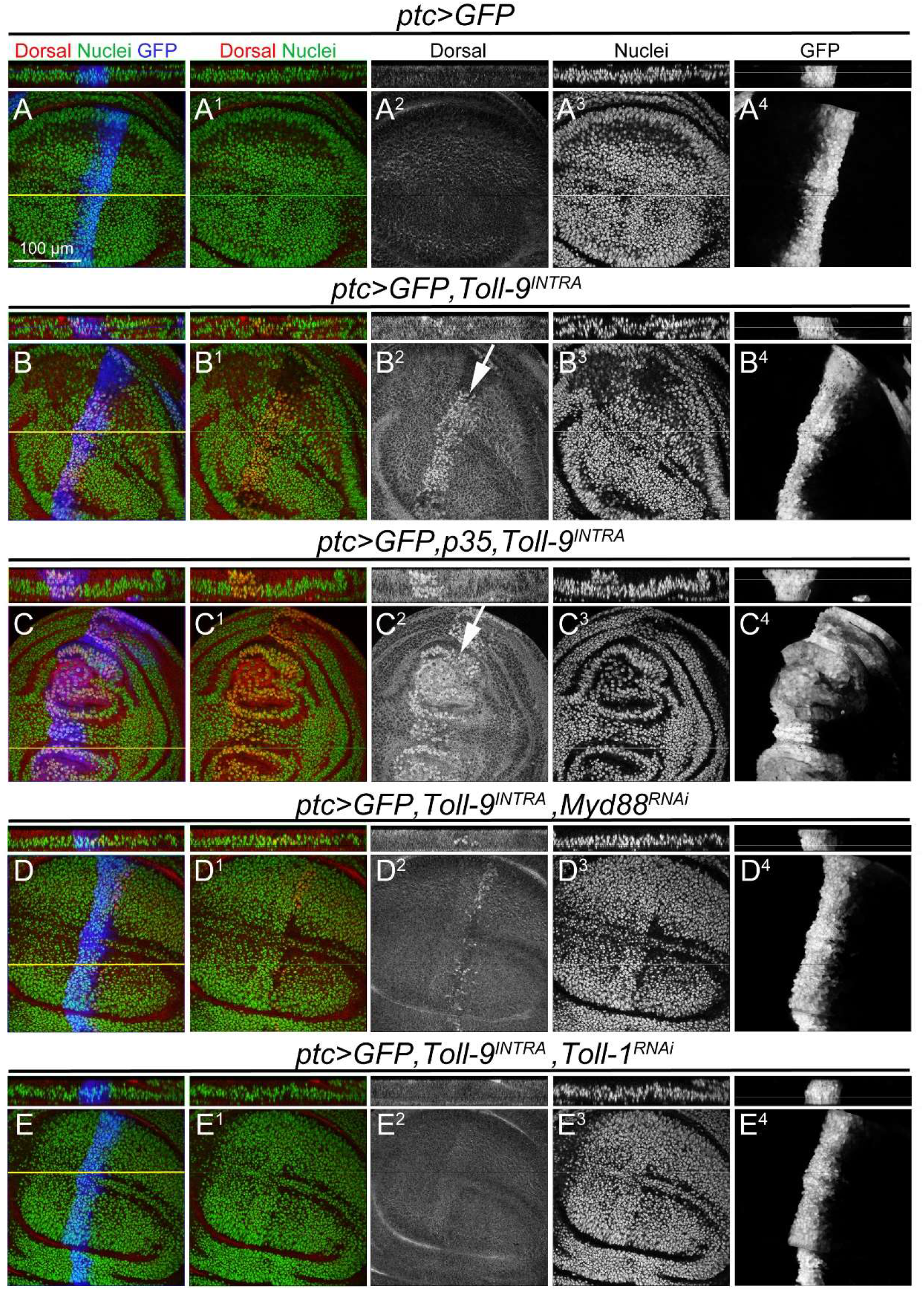
Toll-9-induced nuclear localization of Dorsal is dependent on canonical Toll-1 signaling. All wing imaginal discs were labeled with anti-Dorsal antibody (red). GFP is pseudo-colored in blue and highlights the *ptc* domain. Nuclei (green) were stained with Hoechst. Scale bar: 100μm. The yellow lines in merged panels indicate the regions from which orthogonal sections (shown above panels) were derived. (**A**) A control *(ptc>GFP)* wing imaginal disc. (**B,C**) Misexpression of *Toll-9* results in nuclear localization of Dorsal in *ptc>GFP,Toll-9^INTRA^* (B) and *ptc>GFP,p35, Toll-9^INTRA^* (C) wing imaginal discs. White arrows point to nuclear Dorsal in the *ptc* expression domain. Notably, the *ptc* expression domain was significantly expanded in *ptc>GFP,p35,Toll-9^INTRA^* (C) wing imaginal discs, confirming that in the presence of *p35*, expression of *Toll-9* induces overgrowth. (**D,E**) RNAi knockdown of *Myd88* (D) and *Toll-1* (E) blocks Toll-9-induced nuclear localization of Dorsal.

### Toll-9 laterally interacts with Toll-1 to stimulate nuclear localization of Dorsal

These observations raise the question whether Toll-9 can directly promote the nuclear localization of Dorsal by engaging the Myd88/Tube/Pelle intracellular pathway or whether it laterally interacts with the Toll-1 receptor which then activates the intracellular pathway for nuclear translocation of Dorsal. To address this question, we examined the effect of *Toll-1* RNAi on the *ey>p35, Toll-9* overgrowth phenotype. As shown in Figure 4G (quantified in 4H), *Toll-1* RNAi is a strong suppressor of *ey>p35, Toll-9* induced overgrowth. We also observed that the nuclear localization of Dorsal in the *ptc>p35,Toll-9* expression domain is abolished by *Toll-1* RNAi (Figure 5E). These data imply that Toll-9 can directly or indirectly interact with Toll-1 to engage the known intracellular signaling pathway of Toll-1 for nuclear localization of Dorsal.

### Toll-9 induces of *reaper* and hid in p35-expressing tissue

In the models of cell competition, evidence was provided that Toll-9 contributes to the expression of the pro-apoptotic genes *reaper* and hid, but it was not directly demonstrated that Toll-9 has this activity (Meyer et al., 2014). Consistently, RNAi targeting *reaper* strongly suppressed *ey>p35,Toll-9-induced* overgrowth (Figure 6A,B; quantified in Figure 6D). Furthermore, depletion of *hid* by RNAi also significantly suppresses *ey>p35,Toll-9-induced* overgrowth (Figure 6C,D). These observations suggest that *reaper* and *hid* are required for *Toll-9* induced overgrowth. This notion is further supported by examining the expression of *reaper* and *hid* in *ey>p35,Toll-9* background using *lacZ* reporter genes (Figure 6E-H; quantified in Figure 6I). Interestingly, despite the nuclear localization of Dorsal in *ptc>Toll-9* imaginal discs (Figure 5B), *Toll-9* is only able to induce the expression of the *rpr* and *hid* reporter transgenes in the presence of *p35* (Figure 6F,H,I). The lack of *reaper* and *hid* induction in *ey>Toll-9* discs also explains why there is no significant increase of caspase activity in these discs, while *ey>p35,Toll-9* discs have increased caspase activity (Figure 3C-F) and are dependent on Dronc for overgrowth (Figure 3A,B). Given that expression of *hid* and *reaper* in the presence of *p35* induces overgrowth (see for example Figure 1B), these observations explain why *ey>p35,Toll-9* animals have overgrown heads.

**Figure 6.**
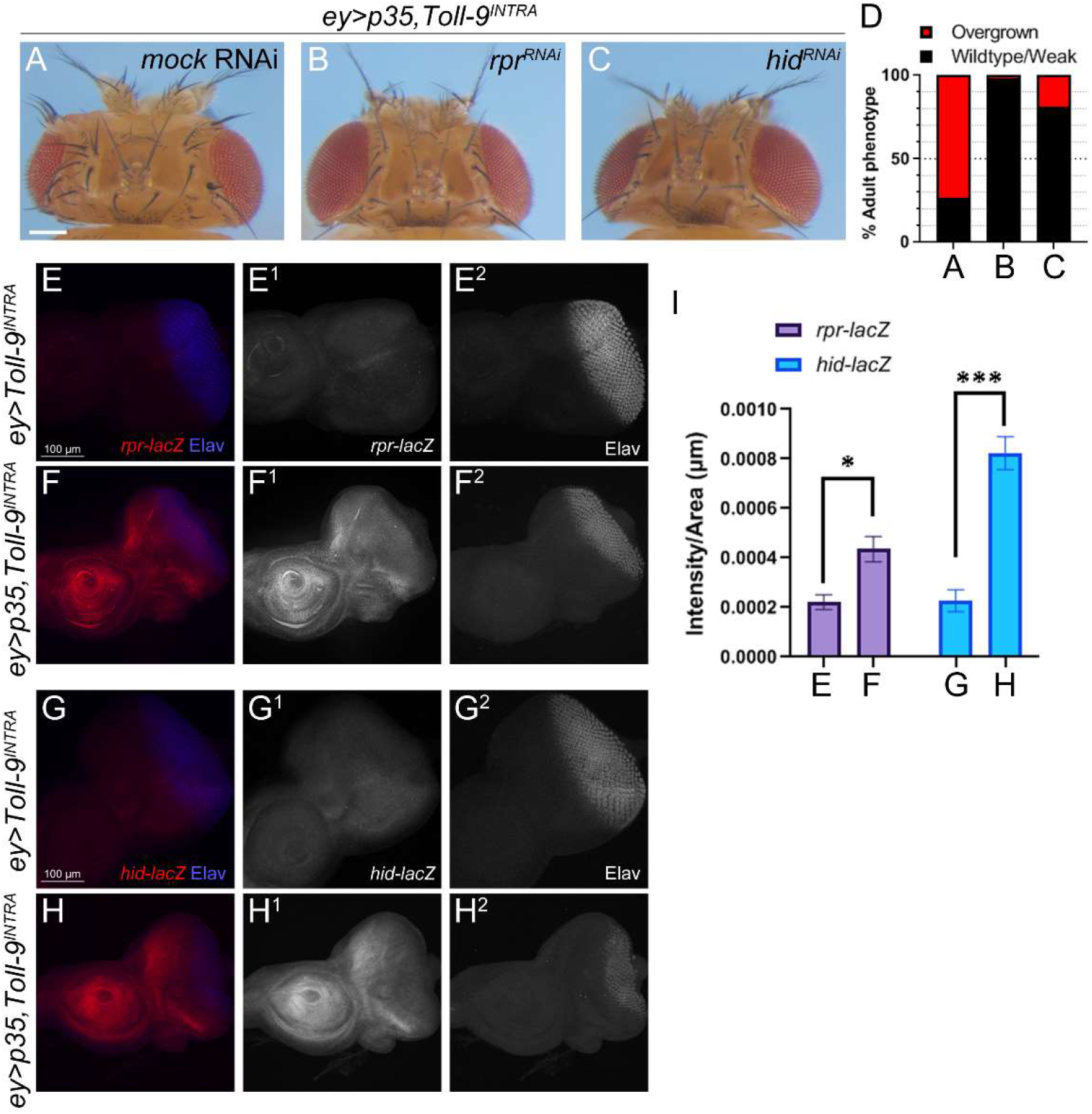
*Toll-9* induces expression of *reaper* and *hid* in *ey>p35* tissue. (**A-C**) Toll-9-induced overgrowth of *ey>p35* head capsules is suppressed by RNAi knockdown of Reaper (B) and Hid (C). Scale bar: 200μm. (**D**) Quantification of the data in panels (A-C). Scoring criteria as in Figure 1E. At least 100 flies per genotype were analyzed. (**E-H**) Toll-9-induced expression of transcriptional reporters of *reaper (rpr-lacZ)* (E,F) and *hid* (*hid-lacZ*) (G,H) in *ey>p35* eye discs. Anti-beta-galactosidase was used to detect *lacZ* expression. Elav labels photoreceptor neurons. Scale bars: 100μm. (**I**) Quantification of the data in panels (E-H). Unpaired t-test, two-tailored distribution, was performed to test the significance. Plotted is the mean intensity ± SEM. ***P<0.001; *P<0.05. The following numbers of discs were analyzed: 7 (E), 15 (F), 4 (G), 17 (H).

## Discussion

In this paper, we examined the role of Toll-9 for AiP because *Toll-9* is the most closely related *Drosophila* TLR compared to mammalian TLRs and a biological function of Toll-9 has not been clearly defined. All mammalian TLRs are involved in innate immunity; therefore, the close homology has led to the prediction that *Drosophila* Toll-9 also participates in innate immunity (Ooi et al., 2002). However, in *Toll-9* mutants, the basal as well as bacterially induced AMP production is not affected, leading to the conclusion that Toll-9 is not involved in innate immunity (Narbonne-Reveau et al., 2011). Nevertheless, instead of eliminating foreign pathogens, previous work has shown that Toll-9 together with several other Toll-related receptors participates in elimination of unfit cells during cell competition (Alpar et al., 2018; Meyer et al., 2014). This is achieved through the expression of the pro-apoptotic gene *reaper*. Here, we demonstrate that Toll-9 has a similar *reaper*- and *hid*-inducing function during AiP, thereby adding to the database of Toll-9 function.

To identify the mechanism by which Toll-9 participates in AiP, we took advantage of the observation that mis-expression of Toll-9 is sufficient to induce overgrowth of *ey>p35* animals. There are multiple aspects of this phenotype which are worth being discussed. First, key for many of the observations presented in this paper is the presence of P35, a very efficient inhibitor of the effector caspases DrICE and Dcp-1. In the absence of P35, overexpression of Toll-9 does not induce any obvious phenotypes (see for example Supplemental Figure S1C). The exact reason for this P35-dependence is currently unknown, but has also been observed upon mis-expression of other genes involved in AiP such as Myo1D (Amcheslavsky et al., 2018). However, because the only known function of P35 is to inhibit DrICE and Dcp-1, it is conceivable that these caspases cleave and inactivate an as-yet unidentified component of the AiP network, possibly to block inappropriate AiP under normal conditions. In the presence of P35, the AiP-blocking activity of DrICE is inhibited and with the addition of an AiP-inducing stimulus such as misexpression of *Toll-9*, AiP is engaged and can trigger tissue overgrowth.

Second, our data show that ectopic *p35, Toll-9* expression triggers overgrowth through a similar mechanism than *hid,p35* expression. This includes Dronc activation, Duox-generated ROS, hemocyte recruitment and JNK activation. These similarities allow us to place the function of Toll-9 into the AiP network.

Third, mis-expressed Toll-9 can genetically interact with Toll-1. This interaction results in nuclear translocation of Dorsal and is dependent on Myd88, Tube and Pelle, all canonical components of the intracellular Toll-1 signaling pathway (Horng and Medzhitov, 2001; Letsou et al., 1991; Shelton and Wasserman, 1993; Tauszig-Delamasure et al., 2002; Towb et al., 2001). Importantly, the nuclear translocation of Dorsal and the *p35,Toll-9* induced overgrowth is also dependent on Toll-1 suggesting that the activation of the Myd88/Tube/Pelle pathway is directly triggered by Toll-1 and not by Toll-9. Mechanistically, Toll-9 may activate Toll-1 either directly through hetero-dimerization or mediated by additional factors. Future work will be necessary to identify the molecular mechanism of the Toll-9/Toll-1 interaction.

Fourth, the outcome of the Toll-9/Toll-1 interaction is the expression of the pro-apoptotic genes *reaper* and *hid*. Because *Toll-9* expression is strongly up-regulated in undead cells (Figure 1F), these data suggest that Toll-9-induced expression of *reaper* and *hid* in undead cells is setting up an amplification loop (Figure 7). The cause of the strong upregulation of Toll-9 in undead cells is unknown. However, other genes are also known to be deregulated in undead cells (Martin et al., 2009). The Toll-9 amplification loop contributes to the strength of undead signaling during AiP and propels the overgrowth of the tissue.

**Figure 7.**
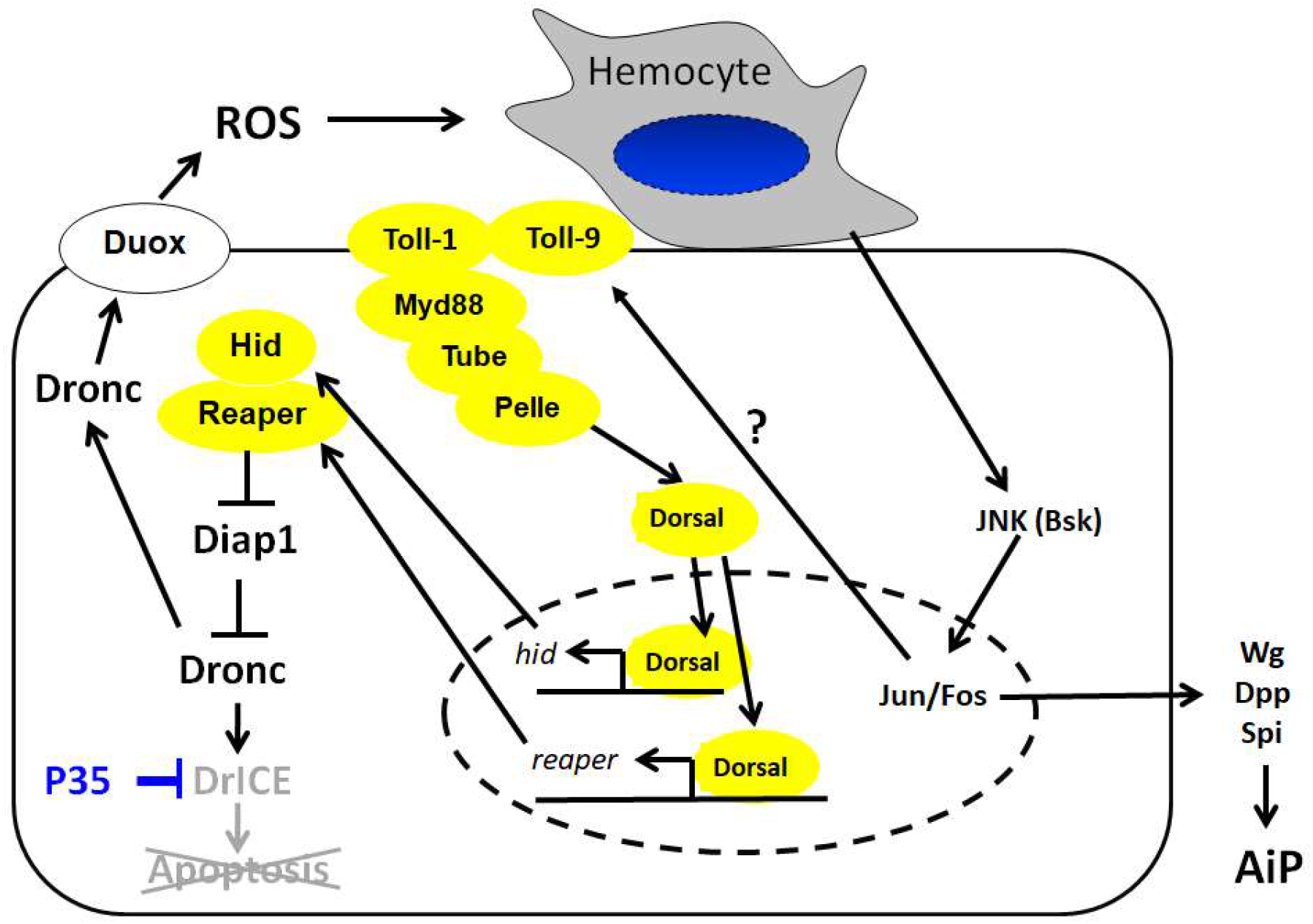
Model of Toll-9 function during apoptosis-induced proliferation (AiP). Shown is an undead epithelial cell of a larval eye-antennal imaginal disc and an attached hemocyte. The interactions revealed in this paper are highlighted in yellow. Toll-9 engages a Toll-1 dependent intracellular pathway culminating in the nuclear translocation of the NF-κB factor Dorsal which directly or indirectly expresses *reaper* and *hid*. Toll-9 may act in a ligand-independent manner. Question marks indicate uncertainty. Previously shown interactions including the activation of Duox, recruitment of hemocytes, and activation of JNK (Amcheslavsky et al., 2018; Fogarty et al., 2016) are also shown. Wg, Dpp and Spi are mitogens produced by undead cells for AiP.

Fifth, another important question is how Toll-9 becomes activated during AiP. Toll-1 requires the ligand Spätzle (Spz) for activation (Morisato and Anderson, 1994; Weber et al., 2003). The *Drosophila* genome encodes *spz* and five additional *spz* homologs (*spz2 – spz6*) (Parker et al., 2001). Targeting all six *spz* genes by RNAi did not suppress *ey>p35,Toll-9-induced* overgrowth (Supplemental Figure S5) suggesting that they do not encode putative Toll-9 ligands. This result is consistent with a recent finding that *Toll-9* RNAi cannot suppress apoptosis induced by a dominant active *spz processing enzyme* (*SPE^Act^*) transgene (Alpar et al., 2018). Although we cannot rule out that there is another unknown ligand, Toll-9 may not need to be activated by a ligand. Toll-9 naturally carries an amino acid substitution in the cysteine-rich extracellular domain similar to the gain-of-function *Toll-1^1^* mutant (Ooi et al., 2002; Schneider et al., 1991). Indeed, Toll-9 behaves as a constitutively active receptor in cell culture assays (Ooi et al., 2002) and the strong transcriptional up-regulation of Toll-9 (Figure 1F) in undead tissue may be sufficient for a detectable response. Since TLRs can form homo- and heterodimers (Hu et al., 2004; Zhang et al., 2002), it is possible that the constitutive activity of Toll-9 might mediate Toll-1 activation in the absence of a natural ligand such as Spz. The strong upregulation of Toll-9 in undead tissue may be the trigger for the Toll-1 activation. Therefore, in a sense, Toll-9 would qualify as a cis-ligand of Toll-1.

With these considerations in mind, the following model for Toll-9 function during undead AiP emerges (Figure 7). The initial stimulus for AiP is the Gal4-induced expression of *hid* and *p35* which leads to the activation of Dronc. Because of P35, Dronc cannot induce apoptosis, but instead activates Duox for generation of ROS (Figure 7). ROS attract hemocytes which release signals for JNK activation in undead cells. Toll-9 is transcriptionally induced. Up-regulated Toll-9 interacts with and activates Toll-1 to engage the Myd88/Tube/Pelle pathway which leads to the nuclear accumulation of Dorsal and potentially Dif (Figure 7). These factors transcriptionally induce *reaper* and *hid* setting up the feedback amplification loop which maintains and propels AiP and overgrowth.

Although *Toll-9* in *Drosophila* does not appear to be required for innate immunity based on its non-essential function to induce AMP production during bacterial infection (Narbonne-Reveau et al., 2011), our work and work by others (Meyer et al., 2014) reveals that Toll-9 may have a function during stress responses which involves expression of pro-apoptotic genes such as *reaper* and *hid*. That was demonstrated previously for cell competition (Meyer et al., 2014) and now for AiP, indicating potential similarities between cell competition and AiP. At first, such similarities appear to be at odds with the common dogmas that proliferating winner cells trigger apoptosis of loser cells, while during AiP, apoptotic cells induce proliferation of surviving cells. However, it has also been reported that loser cells have a much more active role during cell competition and can promote the winner status of cells with increased fitness ((Kucinski et al., 2017). Therefore, there appear to be significant similarities between cell competition and AiP. The common denominator for both systems is the expression of pro-apoptotic genes. However, these responses have different outcomes in both systems. During cell competition, this response involves the death of the loser cells (Alpar et al., 2018; Meyer et al., 2014). During undead AiP, it sets up the amplification loop known to operate in undead cells (Fogarty et al., 2016) which propels hyperplasia and tissue overgrowth.

One other interesting question to examine in the future will be how the intracellular pathway of Toll-1 signaling including Dorsal and Dif can induce different target genes under different conditions. During immune responses, it promotes the expression of AMP genes, while during cell competition and AiP, proapoptotic genes are induced. One potential answer to this question, is that Toll-1 signaling is modified by the interaction with Toll-9. Although this interaction occurs at the level of the plasma membrane, it also appears to influence the activity in the nucleus. It will also be interesting to examine, if Toll-1 can interact with some or all of the other Toll-related proteins in *Drosophila* and how this interaction will influence the specificity of the transcriptional outcome.

## Materials and Methods

### Genetics and fly stocks

All crosses were performed on standard food at 25°C. For expression of RNAi transgenes and mis-expressing transgenes the Gal4/UAS system was used. The exact genotype of *ey>hid,p35* is *UAS-hid* (on X); *ey-Gal4 UAS-p35* (on 2^nd^) and of *ey>p35* it is *ey-Gal4 UAS-p35* (on 2^nd^).

The following fly stocks were used: *ey>hid,p35* and *ey>p35* (Fan et al., 2014); *UAS-Nox* RNAi (line#4) and *UAS-Duox* RNAi (#44) (Ha et al., 2005)(Tanji et al., 2010); *Toll-9^rv17^, UAS-Toll-9, UAS-Toll-9^INTRA^* and *UAS-Toll-9^EXTRA^* (this study); *UAS-Toll-9* RNAi stocks: BL35035, BL36038 and v109635; *UAS-Dif* RNAi: v100537; *UAS-dl* RNAi: BL34938 and v105491;*UAS-Myd88* RNAi: v106198; *tube^3^* mutant (Letsou et al., 1991); *UAS-pelle* RNAi: v103774; *UAS-Toll-1* RNAi: v100078; *UAS-rpr* RNAi: BL51846; *UAS-hid* RNAi: v7912, *UAS-dronc* RNAi: v100424; *UAS-bsk* RNAi (v34138); *UAS-Luciferase* RNAi: BL31603 (used as the mock RNAi); 4kb *rpr-lacZ* and *hid[10-1kb]-lacZ* (line1) (Fan et al., 2010); *TRE-RFP* (Chatterjee and Bohmann, 2012); *UAS-spz* RNAi: BL28538; *UAS-spz2* RNAi: v26115; *UAS-spz3* RNAi: BL56958; *UAS-spz4* RNAi: BL60044; *UAS-spz5* RNAi: v102389; *UAS-spz6* RNAi: BL57510.

### Generation of the *Toll-9^rv17^* allele

The parental P element insertion strain EY14405 from Bloomington stock center was used for a jump out excision screen. The strain was crossed with a Δ2-3 strain and offspring with loss of red eye color were collected. Individual flies collected were crossed with balancer strain to establish individual stocks. Genomic DNA from these collected stocks were isolated and tested by PCR using primers surrounding the P element insertion site. The *revertant 17* (*rv17*) stock showed absence of PCR product on primary screen and the homozygous strain were subject to further PCR and sequencing to confirm the deletion of the *Toll-9* locus, from the insertion site in the second intron to the last coding exon as shown in Supplemental Figure S1A.

### Generation of *UAS-Toll-9^INTRA^* and *UAS-Toll-9^EXTRA^*

For *<u>UAS-Toll-9^EXTRA^</u>*, the *Toll-9* cDNA was used to PCR amplify the first part of the coding sequence, from 5’ UTR to the beginning of cytoplasmic domain including the transmembrane domain. The clone encodes a.a. 1 to 700. The PCR fragment was cloned into the XbaI and BglII sites of the pUAST vector. For *<u>UAS-Toll-9^INTRA^</u>*, the *Toll-9* cDNA was used to amplify a short fragment encoding the first 15 a.a., as signal peptide sequence, and a second fragment encoding a.a. 650, to the stop codon, including the transmembrane and the cytoplasmic domains. The two fragments were cloned head to tail into the XbaI and BglII sites of the pUAST vector. The resulting plasmids were used to generate transgenic flies (Rainbow Transgene, California) in the *w^1118^* background. The resulting transgenic flies were used for crosses into the *ey>p35* background.

### Immunolabeling and ROS staining

For labelings with antibodies, fixed eye-antennal or wing imaginal discs from 3^rd^ instar larvae were used following standard protocols (Fogarty et al., 2016). The monoclonal NimC antibody is a kind gift of István Andó (Kurucz et al., 2007) and was used at a dilution of 1:300. Primary antibodies from the Developmental Studies Hybridoma Bank (DSHB) included anti-Dorsal (7A4) used at 1:20, anti-Mmp1 (3A6B4) used at 1:50, anti-Wingless (4D4) used at 1:50, anti-beta-galactosidase (40-1a) used at 1:20, and anti-Elav (7E8A10) used at 1:50. Anti-cDcp1 (Asp216) antibody from Cell Signaling Technology was used at 1:200. Hoecst 33342 solution was used at 1:1000 to stain nuclei. Fluorescent secondary antibodies were obtained from Jackson ImmunoResearch. ROS staining with DHE (Invitrogen #D23107) and H_2_DCF-DA (Molecular Probes #C6827) were performed on unfixed imaginal discs using published protocols (Owusu-Ansah et al., 2008).

### Quantification and statistics

For quantification of confocal images, the ‘Histo’ function of Zen 2012 imaging software (Carl Zeiss) was used. Region of interest was outlined for each disc and mean fluorescence signal intensity was determined. Crosses were repeated at least three times. Analysis and graph generation was done using GraphPad Prism 7.04. The statistical method used was one-way ANOVA with Holm-Sidak test for multiple comparisons, unless otherwise indicated. Plotted is mean intensity ± SEM. Levels of significance are depicted by asterisks in the figures: * P<0.05; ** P<0.01; *** P<0.001; **** P <0.0001. Total number of discs used for quantification are indicated in the legends for each figure.

### qRT-PCR for determination of *Toll-9* transcript levels

Total mRNA was prepared from eye-imaginal discs using TRIzol reagent (Ambion, Cat#15596026). cDNA was prepared using QuantiTect Reverse Transcription Kit (Qiagen, Cat# 205311) and qPCR was performed with Power SYBR Green PCR master mix (Applied Biosystems, Cat#4367659). The following primer pairs were used:

Toll-9, Forward: 5’-CCATTACAAGCACTATAGG;

Reverse: 5’-GACCTCTTCGGCCTCTTC.

RP-49: Forward: 5’-CCAGTCGGATCGATATGCTAA;

Reverse: 5’-ACGTTGTGCACCAGGAACTT.

## Acknowledgments

We would like to thank István Andó, Dirk Bohmann, Won Jae Lee, the Bloomington *Drosophila* Stock Center and the Vienna *Drosophila* Resource Center for fly stocks. This work was funded by the National Institute of General Medical Science (NIGMS) under award number R35GM118330 to AB. YTI was supported by NIH grants GM107457 and DK083450. The content is solely the responsibility of the authors and does not necessarily represent the official views of the NIH. YTI is a member of the UMass Center for Clinical and Translational Science (UL1TR000161).

## Author contributions

AS, AA and EB performed most of the experiments shown in the figures; YN generated and assayed the activity of the transgenic *Toll-9*; TT assayed various *Toll-9* deletion constructs and generated the *Toll-9^rv17^* mutant. YTI designed and supervised the generation of *Toll-9* constructs and *Toll-9^rv17^* mutant. AB supervised the experimental part of the project. AS, AA, YTI and AB interpreted the results and wrote the manuscript. AB and YTI acquired funding.

## Data and materials availability

All data is available in the main text or the supplementary materials. All materials are available upon request from the corresponding author.

## List of Supplemental Materials

Figures S1 to S5

## Supplemental Materials

**Supplemental Figure S1.**
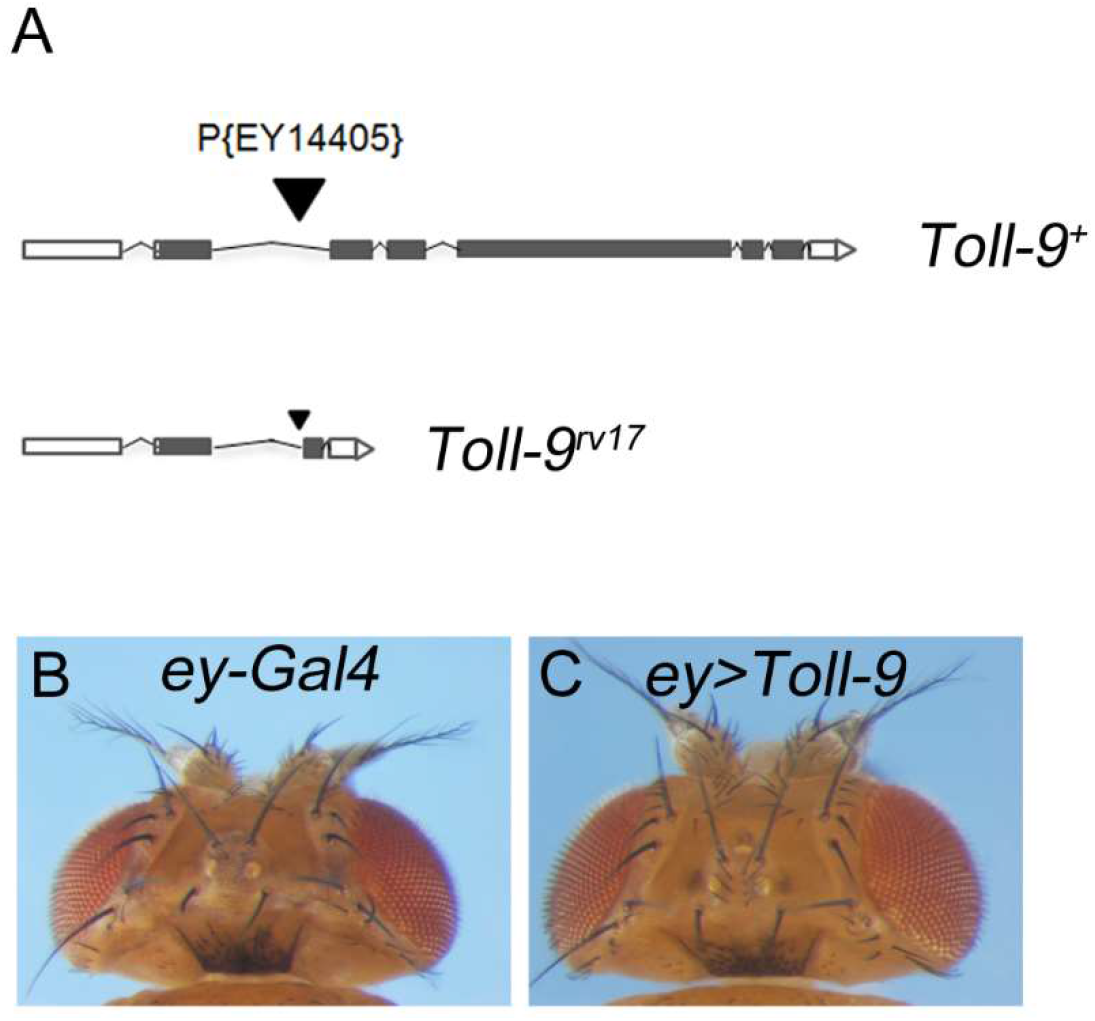
Genomic region of *Toll-9^rv17^* and phenotype of *ey>Toll-9*. (**A**) Outline of the genomic regions of *Toll-9^wt^* and *Toll-9^rv17^*. Specifics about the generation of *Toll-9^rv17^* are described in the Materials and Methods. (**B,C**) Expression of *Toll-9* using *ey-Gal4* does not induce an obvious eye/head phenotype.

**Supplemental Figure S2.**
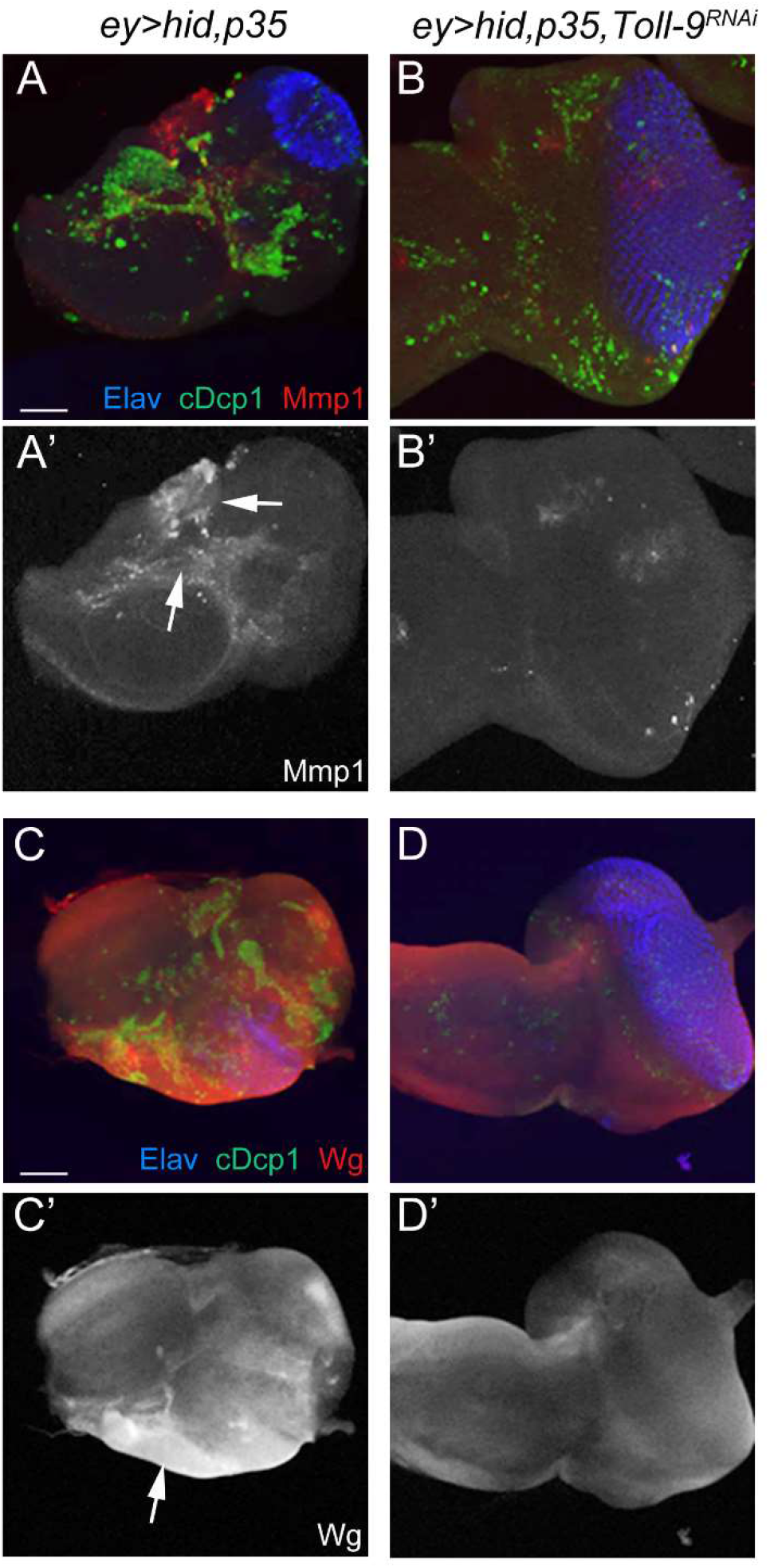
*Toll-9* functions upstream of JNK and Wingless activity in *ey>hid,p35* eye discs. (**A-D’**) JNK activity (A-B’) and Wingless (Wg) expression (C-D’) in *ey>hid,p35* eye imaginal discs were reduced by *Toll-9* RNAi. Arrows in (A’) and (C’) mark examples of strong JNK and Wg activity in ey>hid,p35 discs. Anti-Mmp1 antibody was used as a readout for JNK signaling. Anti-Wg antibody revealed Wingless activity. Elav marked photoreceptor neurons. cDcp1 highlighted cells with active effector caspases. Scale bar: 50μm.

**Supplemental Figure S3.**
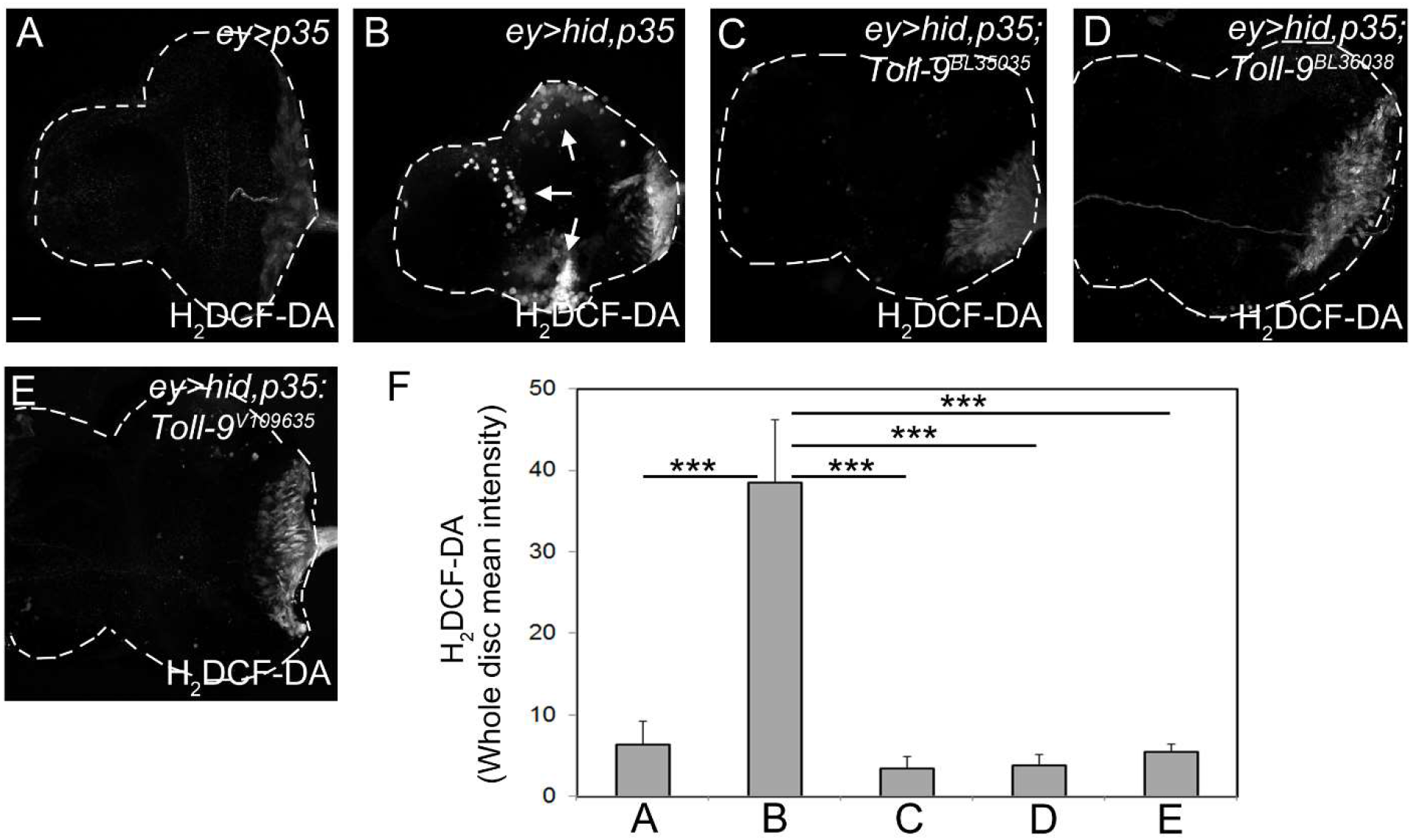
*Toll-9* is required for ROS generation in *ey>hid,p35* discs. (**A-E**) H_2_DCF-DA labeling as ROS indicator of eye imaginal discs of indicated genotype. *Toll-9* knockdown suppresses H_2_DCF-DA labeling (C-E). BL35035, BL36038 and v109635 are the Bloomington and Vienna stock center numbers for the *Toll-9* RNAi stocks used. Arrows in (B) highlights the H_2_DCF-DA labeling in *ey>hid,p35* discs. (**F**) Quantification of (A-E). H_2_DCF-DA labeling was measured across entire discs and normalized to disc size. Data were analyzed by one-way ANOVA with Holm-Sidak test for multiple comparisons. Mean signal intensity is plotted ± SEM. ***P<0.001.

**Figure S4.**
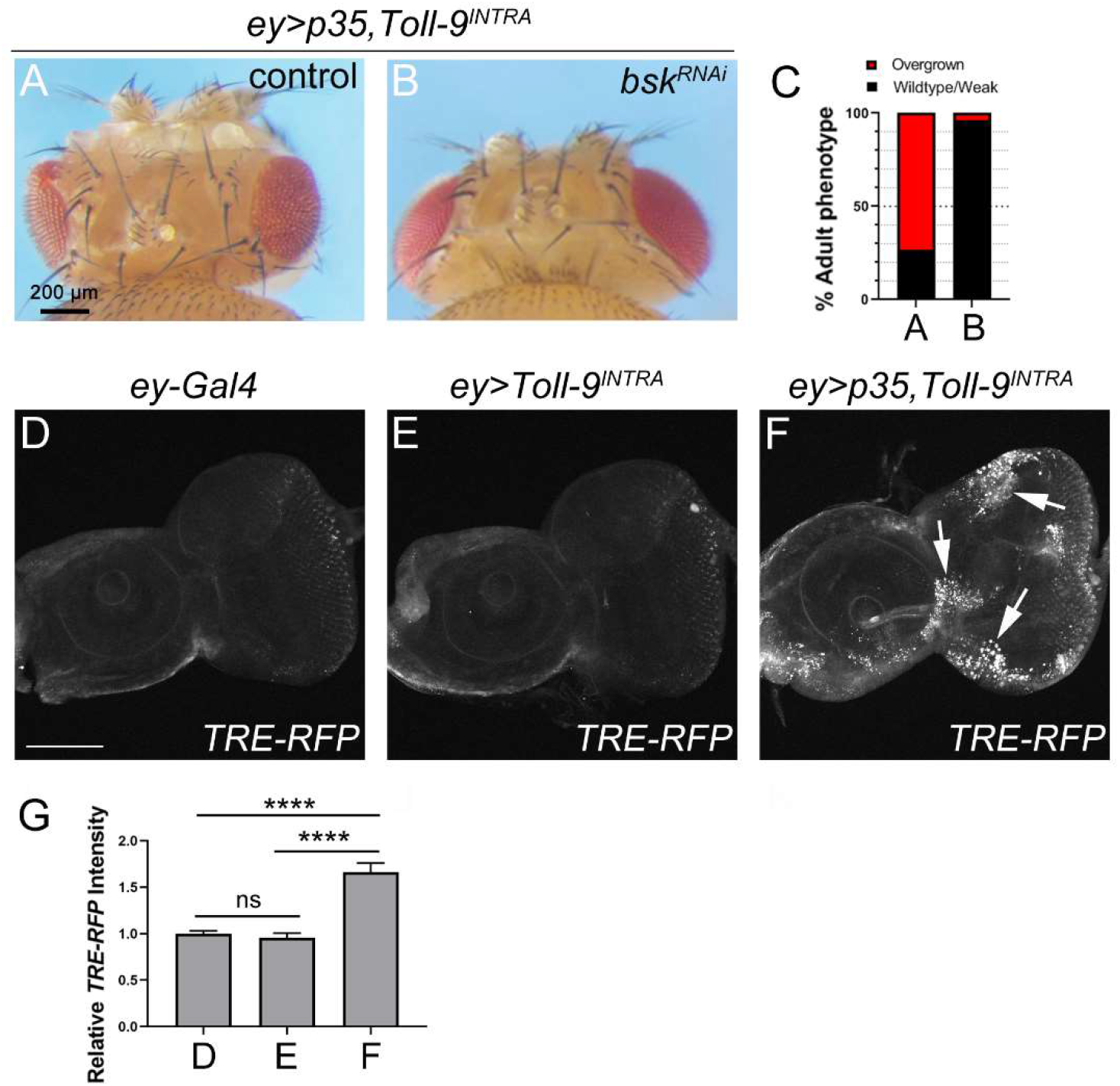
JNK is required for *ey>p35,Toll-9-induced* overgrowth. (**A,B**) RNAi-knockdown of *JNK* encoded by the *basket (bsk)* gene (B) suppressed *ey>p35, Toll-9-induced* overgrowth (A). Scale bar: 200μm. (**C**) Quantification of the data in panels (A,B). Scoring criteria as in Figure 1E. (**D-F**) JNK signaling detected by *TRE-RFP* reporter expression in *ey-Gal4* (D), *ey>Toll-9^INTRA^* (E) and *ey>p35, Toll-9^INTRA^* (F) eye discs. Arrows in (F) highlight a few examples of increased *TRE-RFP* expression. Scale bar: 100μm. (**G**) Quantification of the *TRE-RFP* signals in (D-F). *TRE-RFP* labeling was measured across entire discs and normalized to disc size. Data were analyzed by one-way ANOVA with Holm-Sidak test for multiple comparisons. Mean signal intensity is plotted ± SEM. ****P<0.0001. ns = not significant. n = 8 (D), 9 (E), 12 (F).

**Supplemental Figure S5.**
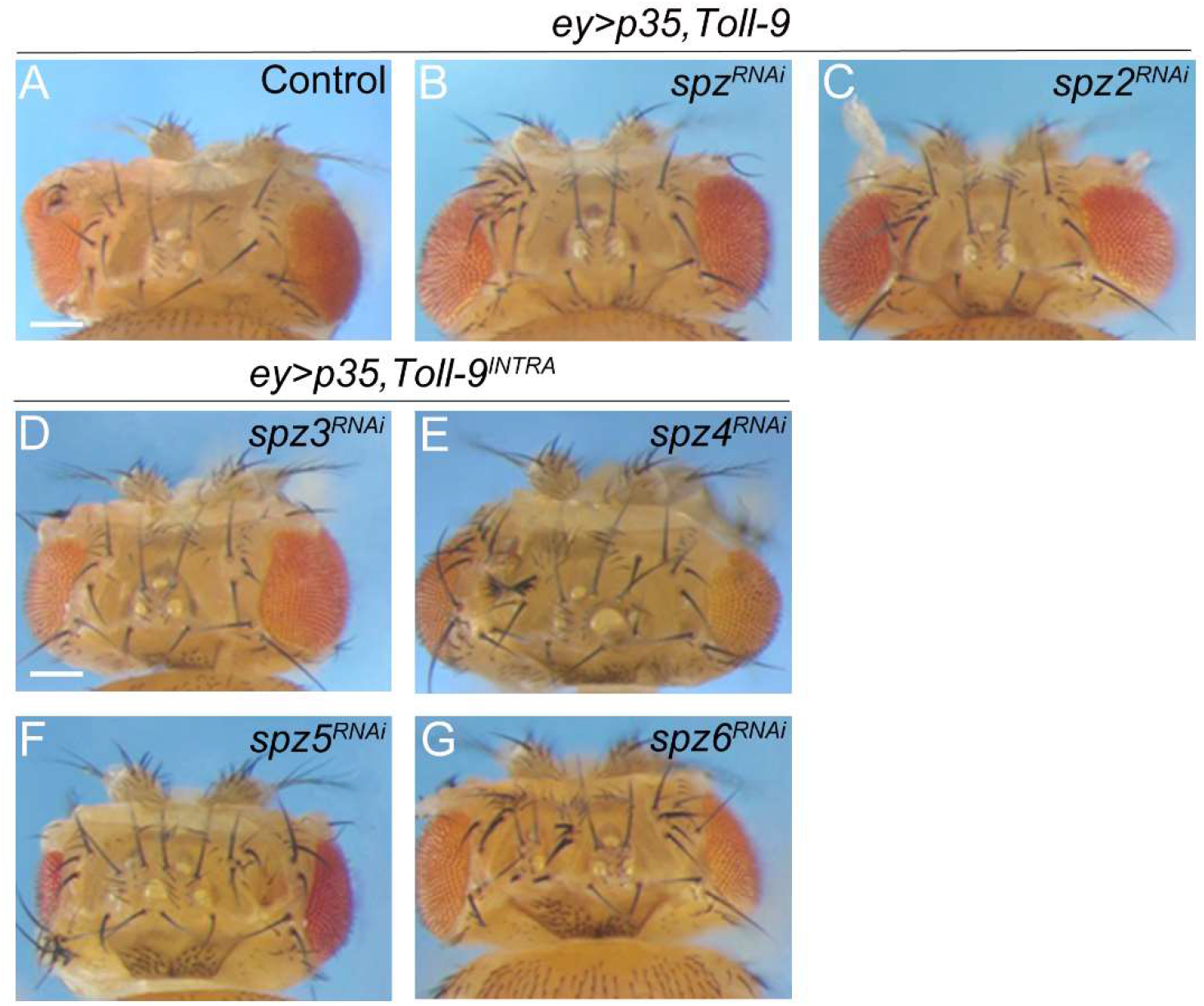
The extracellular Spätzle ligands are not required for *ey>p35*, Toll-9-induced overgrowth. (**A-G**) *ey>p35, Toll-9-induced* overgrowth (A) is not affected by RNAi-mediated knockdown of the six *späetzle* genes (B-G) encoded in the *Drosophila* genome. Scale bars: 200μm.

